# The draft genome of the microscopic *Nemertoderma westbladi* sheds light on the evolution of Acoelomorpha genomes

**DOI:** 10.1101/2023.06.28.546832

**Authors:** Samuel Abalde, Christian Tellgren-Roth, Julia Heintz, Olga Vinnere Pettersson, Ulf Jondelius

## Abstract

**Background:** Xenacoelomorpha is a marine phylum of microscopic worms that is an important model system for understanding the evolution of key bilaterian novelties, such as the nervous or excretory systems. Nevertheless, Xenacoelomorpha genomics has been restricted to the few species that either can be cultured in the lab or are centimetres long. Thus far, no genomes are available for Nemertodermatida, one of the phylum’s main clades and whose origin has been dated more than 400 million years ago.

**Results:** We present the first nemertodermatid genome sequenced from a single specimen of *Nemertoderma westbladi*. Although genome contiguity remains challenging (N50: 48 kbps), it is very complete (BUSCO: 81.4%, Metazoa; 91.8%, Eukaryota) and the quality of the annotation allows fine-detail analyses of genome evolution. Acoelomorph genomes seem to be conserved in terms of the percentage of repeats, number of genes, number of exons per gene and intron size. In addition, a high fraction of genes present in both protostomes and deuterostomes are absent in Acoelomorpha. Interestingly, we show that all genes related to the excretory system are present in Xenacoelomorpha but *Osr*, a key element in the development of these organs and whose acquisition might explain the origin of the specialised excretory system.

**Conclusions:** Overall, these analyses highlight the potential of the Ultra-Low Input DNA protocol and HiFi to generate high-quality genomes from single animals, even for relatively large genomes, making it a feasible option for sequencing challenging taxa, which will be an exciting resource for comparative genomics analyses.

## 1. Background

Access to a growing number of high-quality genomes from non-model animal species has helped us understand the origin of key evolutionary novelties [1–3]. However, small yields of extracted DNA is a limiting factor in genome sequencing of small animals, also when using whole-body extractions. In this regard, the recent development of the Ultra-Low Input DNA protocol has significantly reduced the amount of input DNA, enabling the sequencing of high-quality genomes from millimetric animals [4–6]. Yet, this approach is recommended for genomes smaller than 500 Mbps, and it is unclear how well it performs beyond that limit, which is not a minor detail. Despite the general trend that miniaturised animals tend to have smaller genomes [7–9], there are several phyla, such as Xenacoelomorpha, whose genome size is comparable to that of larger animals [10–12].

Xenacoelomorpha is a phylum of marine, microscopic worms consisting of the clades Acoela, Nemertodermatida, and their sister taxon *Xenoturbella*. Early molecular phylogenetic studies placed Xenacoelomorpha as the sister group of all other Bilateria. This hypothesis received support from the simple morphology of Xenacoelomorpha, which lack typical bilaterian structures such as excretory organs, through-gut and circulatory system [13] and the name Nephrozoa was introduced for its sister group under this hypothesis [14]. The Nephrozoa hypothesis was further supported by analyses of gene content and phylogenomic inference [15,16]. However, an alternative hypothesis based on analyses of nucleotide sequence data places Xenacoelomorpha as sister group to Ambulacraria (echinoderms and hemichordates) within the deuterostomes [17,18]. In either case, xenacoelomorphs offer a good opportunity for studying the origin of important animal novelties. Due to their lack of specialised excretory organs, xenacoelomorphs make a good comparison reference to better understand the evolution of this system. A recent study based on spatial transcriptomics has shown the expression in Xenacoelomorpha of several genes involved in the excretory process in other bilaterians, as well as several genes specifically related to the ultrafiltration excretory system (*Nephrin*, *Kirrel*, and *ZO1*; [19]), although their expression was observed throughout the body, unlike in other organisms with specialised excretory organs [20]. In addition to analysing their expression, the comparison of high-quality genomes from xenacoelomorphs, protostomes, and deuterostomes would offer a better understanding of the evolution of these genes, thanks to a more accurate assessment of gene presence/absence, the annotation of all gene copies in the genome, information about their distribution in the genomes, or comparisons of gene architecture, among other analyses. However, the set of available xenacoelomorph genomes is still limiting.

Several xenacoelomorph species have drawn interest as a model system to study the evolution of body regeneration, the nervous system, and endosymbiosis [12,19,21], resulting in the generation of genomes from *Xenoturbella* (*Xenoturbella bocki*; [22]) and Acoela (*Hofstenia miamia* and the closely related acoel species *Praesagittifera naikaiensis* and *Symsagittifera roscoffensis*; [10–12]). Thus, to fully capture the diversity of Xenacoelomorpha it is necessary to generate new genomes from Nemertodermatida, the sister group of Acoela and from which diverged more than 400 MYBP [23]. This, however, is challenging due to their microscopic size. The four available xenacoelomorph genomes were sequenced from species that can be either cultured in the lab and/or are relatively big (*Xenoturbella* and *Hofstenia* can reach four and two cm body length, respectively), but that is not the case for the vast majority of xenacoelomorphs, requiring more sophisticated methods. Despite their small size, all acoel genomes sequenced so far range between 700 and 1000 Mbps, two to three times larger than any other genome sequenced with the Ultra-Low Input protocol so far [4–6], and thus represent a good opportunity for testing its performance in a challenging animal group. Here, we applied the PacBio Ultra-Low DNA Input protocol to sequence the genome of *Nemertoderma westbladi* from a single, microscopic worm, the first nemertodermatid and the longest genome sequenced with this protocol. We demonstrate the potential of this approach to generate relatively good-quality genomes through comparisons with other genomes from this phylum. In addition, we explore the evolution of acoelomorph genomes, analyze the evolution of gene content in Bilateria and provide insights into the evolution of the genes related to the excretory system.

## 2. Results

### 2.1. The *Nemertoderma westbladi* genome

The best extraction was produced from a sample stored in RNAlater using the QIAamp Micro kit, obtaining a fragment size over 20 kbps and ca. 20 ng of total DNA, which would be up to 990 ng after DNA shearing and whole genome amplification. About half of this DNA was selected for sequencing. A total of 2,313,071 reads were produced during HiFi sequencing, later reduced to 2,297,478 after quality filtering with an average length of 6.6 kbps.

Flye produced the best assembly, which was 678.9 Mbps long and contained 26,880 contigs (Fig. 1A). The longest contig was 2 Mbps long, with an N50 of 42.6 kbps and contained 86.6% of the BUSCO Metazoa odb10. The assembly contained two repeats of 507 and 531 bps with 70,000 and 79,000 copies, respectively, corresponding to 11% of the assembled genome. BlobTools2 revealed the presence of many contaminants, with only 61% of the contigs identified as metazoan (Supplementary Table S1). Thus, the decontaminated assembly was only 558.6 Mbps, split into 15,300 contigs with an N50 of 48.17 kbps (Fig. 1A, Table 1), but 81.4% of the Metazoa and 91.8% of the Eukaryota BUSCO genes were still present (Supplementary Figure S1). The smudgeplot was markedly different before and after the decontamination step, as the inferred ploidy went from triploid to diploid after the decontamination (Supplementary Figure S2). The genome size estimated by GenomeScope was 235.4 Mbps, with an average coverage of 24.3, and high heterozygosity (6.45%), although these numbers must be taken cautiously given the poor fit of the model (33%; Supplementary Figure S3).

**Figure 1:**
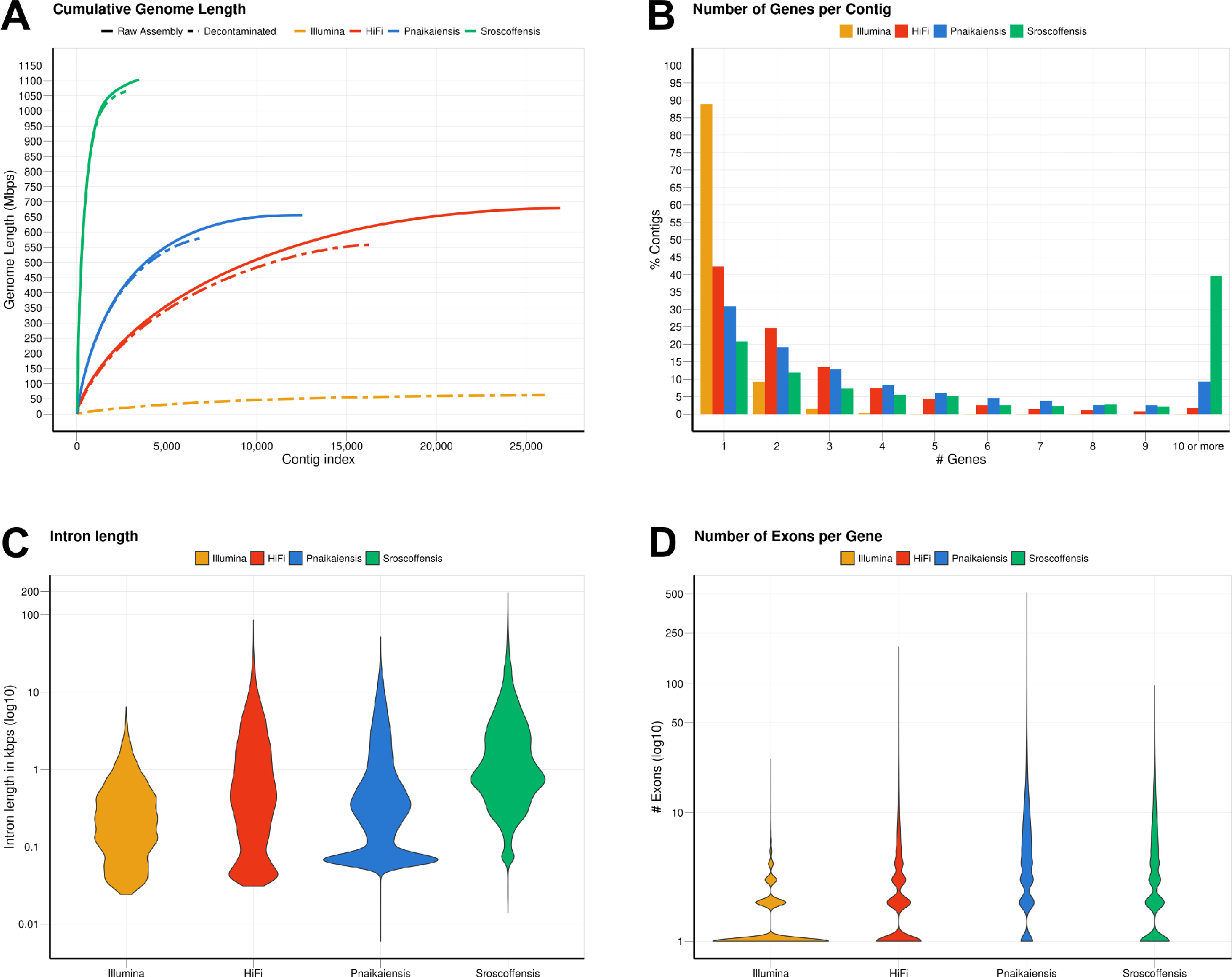
Summary of the statistics calculated for the two *N. westbladi* genomes (sequenced with Illumina or HiFi), *P. naikaiensis*, and *S. roscoffensis*. (A) Cumulative genome length, sorted from the longest to the shortest contig, separating the raw assembly from the BlobTools decontamination. Due to the large number of contigs in the raw assembly, only the decontaminated version of the *N. westbladi* genome sequenced with Illumina is shown. (B) Summary of the number of genes per contig, (C) distribution of the intron length per species, and (D) number of exons per gene.

**Table 1:** Statistics of the four genomes analysed in this study after the decontamination step. The *N. westbladi* genomes are presented as “HiFi” and “Illumina” to differentiate the two sequencing approaches.

| Parameter | Illumina | HiFi | Pnaikaiensis | Sroscoffensis |
| --- | --- | --- | --- | --- |
| Length after BlobTools (Mbps) | 62.229 | 558.589 | 581.371 | 1064.926 |
| N's (count) | 49,310 | 15,300 | 7,367,142 | 1,589,933 |
| N's (%) | 0.079 | 0.003 | 1.267 | 0.149 |
| Number of contigs | 26,021 | 16,265 | 7104 | 2730 |
| Longest contig (Kbps) | 65.353 | 601.587 | 702.461 | 8003.794 |
| Average contig length (Kbps) | 2.391 | 34.343 | 81.837 | 390.083 |
| N50 (Kbps) | 3.996 | 48.170 | 129.752 | 1077.644 |
| Number of gene models | 23,120 | 30,698 | 20,303 | 28,513 |
| Fuctionally annotated proteins | 14,486 | 12,849 | 13,708 | 17,717 |
| Max. number of genes per contig | 33 | 89 | 37 | 280 |
| Average number of genes per contig | 0.876 | 1.816 | 2.858 | 12.281 |
| Max. number of exons per gene | 26 | 195 | 512 | 97 |
| Average number of exons per gene | 1.531 | 3.044 | 6.386 | 4.244 |

The decontaminated Illumina genome was also relatively complete, with 76.8% of the metazoan BUSCO genes present in the assembly, but much shorter (62.2 Mbps) and much more fragmented (49,310 contigs; N50: 4 kbps) (Fig. 1A). Despite being sequenced from cultured, starved and free of symbionts populations, BlobTools also identified some contaminants in the published genomes of *P. naikaiensis* and *S. roscoffensis*. The former went from 656.1 Mbps and 12,525 contigs to 581.4 Mbps and 7104 contigs, whereas the latter went from 1103 Mbps and 3460 contigs to 1064.9 Mbps and 2730 contigs (Fig. 1A). The N50 of the two genomes raised from 127 to 130 kbps in *P. naikaiensis*, and from 1.04 to 1.08 Mbps in *S. roscoffensis*. Despite the observed differences in genome size and contiguity, the four genomes show very similar completeness results. More than 90% of the Eukaryota BUSCO genes were identified in the decontaminated genomes of all species but *P. naikaiensis* (14.9% of missing genes) (Supplementary Figure S2A). Differences were slightly higher with the Metazoa database, with almost a 10% difference between the most (*S. roscoffensis*; 18.5% missing genes) and the least (*P. naikaiensis*; 27%) complete genomes. In *N. westbladi*, the HiFi genome was almost as complete as *S. roscoffensis* (18.6% missing genes), whereas the Illumina genome was in an intermediate position (23.1%) (Supplementary Figure S2B).

The number of gene models in the four genomes ranged from 20,303 (*P. naikaiensis*) to 30,698 (*N. westbladi*, HiFi genome), although the differences were reduced when only functionally annotated genes were considered: 12,849 (*N. westbladi*, HiFi), 13,708 (*P. naikaiensis*), 14,486 (*N. westbladi*, Illumina), and 17,717 (*S. roscoffensis*) (Table 1). The organisation of these genes in the genome somehow reflected the differences observed in genome contiguity. In the *N. westbladi* genome sequenced with Illumina, the average number of genes per contig was just 0.876, with a single gene in almost 90% of the contigs (Fig. 1B), and the contig with the highest number of genes presented 33 gene models (Table 1). In the HiFi sequenced *N. westbladi* genome, up to 89 genes were found in a single contig, with an average of 1.8 genes per contig. Similarly, an average of 2.9 genes per contig were annotated in the *P. naikaiensis* genome, but in this case, the maximum number of genes in one contig was only 37. The *S. roscoffensis* genome stands out, with a maximum of 280 genes in a single contig and more than 10 genes in almost 40% of the contigs (Fig. 1B; Table 1). This trend, however, was not observed in gene architecture. The gene models in *P. naikaiensis*, *S. roscoffensis*, and the HiFi genome of *N. westbladi* were similar, ranging between an average of 3 to 6.3 exons per gene, whereas almost all the genes presented a single exon in the Illumina genome (average 1.5) (Fig. 1D). The intron size was very variable in all genomes, ranging from 6 (*P. naikaiensis*) to 193,733 (*S. roscoffensis*) bps. The intron size distribution was similar between *N. westbladi* and *P. naikaiensis*, but with generally longer introns in *S. roscoffensis* (Fig. 1C). Nevertheless, the intron size range was similar in the three genomes, but visibly smaller in the *N. westbladi* Illumina genome.

According to RepeatMasker, the *N. westbladi* genome is very repetitive, masking up to 59.85% of the genome (Supplementary Table S2). The majority of these repeats are interspersed throughout the genome (58.34%) and more than a fifth (21.27%) were not classified into any known repeat family. Among the classified repeats, the most common ones are retroelements (33.36%), particularly the long terminal repeats (LTR, 21.87%) and long interspersed nuclear elements (LINEs, 11.15%). The Illumina genome presents a sharp contrast, with just 16.40% of the genome masked as repetitive, although LINEs (4.28%) and LTR (3.43%) are still the most abundant repeat elements (Supplementary Table S2).

### 2.2. Identification of the contaminant contigs

More than half of the taxonomic groups identified within the set of contaminant contigs were bacteria, including several of the major taxonomic groups: Bacteroidetes, Tectomicrobia, Proteobacteria (including Alpha-, Beta-, Delta/Epsilon-, and Gammaproteobacteria), Planctomycetes, Actinobacteria, Cyanobacteria, and Firmicutes. None of the “Candidate Phyla Radiation” phyla were identified. More specifically, there are nine genera that have been reported as statistically more abundant in the microbiome of microscopic animals than in environmental samples [24] and thus might be part of the *Nemertoderma* microbiome: *Algoriphagus*, *Alteromonas*, *Francisella*, *Photobacterium*, *Roseobacter*, *Shewanella*, *Streptococcus*, *Tenacibaculum*, and *Vibrio*. Other important sources of contamination besides Bacteria are algae (Chlorophyta, Rhodophyta, and Streptophyta), land plants (Streptophyta: Bryopsida and Spermatophyta), and fungi (Ascomycota, Basidiomycota, Chytridiomycota, Microsporidia, Mucoromycota, and Zoopagomycota). These groups accumulate 87% of the taxonomic diversity within the contaminants. In addition, we also found Archaea (Thaumarchaeota: *Nitrososphaera*), Protista (Amoebozoa, Euglenozoa, Apicomplexa, Ciliophora, Perkinsozoa, Endomyxa, and Oomycota), and Virus (Uroviricota and Nucleocytoviricota). A complete description of these results is provided in Supplementary Table S3.

### 2.3. Gene content evolution

The comparison of 18 animal genomes, representing Acoelomorpha, Cnidaria, Deuterostomia, and Protostomia revealed a high degree of specificity in gene content: 17.4% of all orthogroups present in Cnidaria are exclusive to this phylum, 24.6% in Acoelomorpha, 45.4% in Deuterostomia, and 48.6% in Protostomia (Fig. 2A). Hence, only 35.9% of all orthogroups were annotated in at least two of the four groups (12,071 out of 33,649). Among these, almost half (47.6%) were present in at least one species of each clade, whereas only 3.4% were present in all bilaterian clades but Cnidaria. A total of 8,394 genes were identified as shared across Metazoa (present in Cnidaria and at least one Bilateria), and 2,328 for Bilateria (present in at least two bilaterian clades). Acoelomorpha was present in 71.8% of the metazoan genes and 42.1% of the bilaterian ones, contrasting with deuterostomes (91.4% and 82.9%) and protostomes (94.5% and 92.6%) (Fig. 2C). The proportion of missing BUSCO genes was below 11% in all four groups (Fig. 2B), and so genome completeness does not explain this pattern. Within Acoelomorpha, almost half (43.8%) of the genes were shared between Acoela and Nemertodermatida (Fig. 2A).

**Figure 2:**
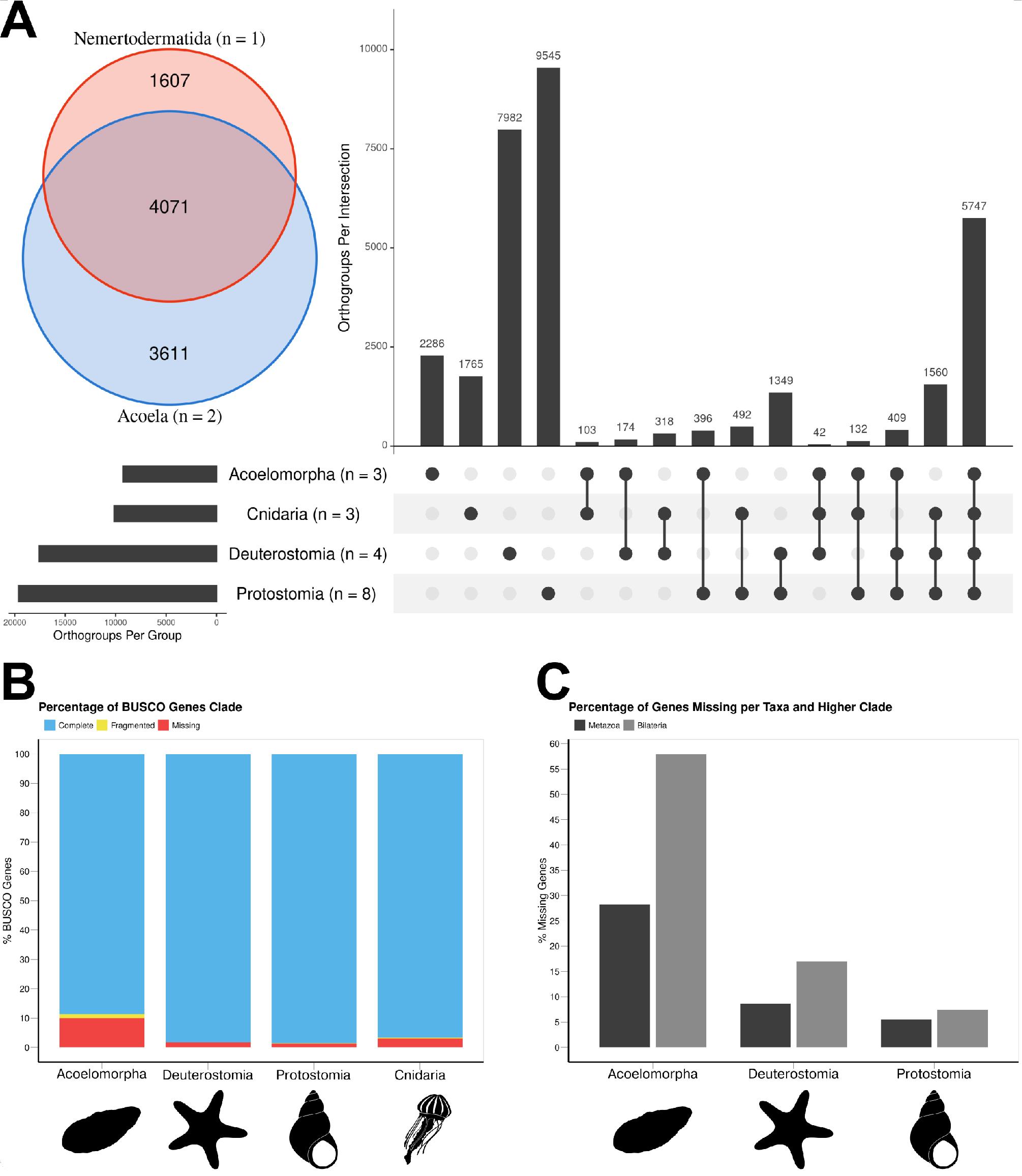
The gene content of the three acoelomorph genomes was compared to 15 genomes from several phyla, including three cnidarians, four deuterostomes (three chordates and one echinoderm), and eight protostomes. (A) Number of unique and shared genes among acoelomorphs, cnidarians, deuterostomes, and protostomes. In the inset, the number of shared genes between the two acoel genomes and *N. westbladi*. (B) BUSCO scores of each of the four main clades. (C) Percentage of missing genes observed in acoelomorphs, deuterostomes, and protostomes. The set of “metazoan genes” was defined as all genes shared between at least one cnidarian and one bilaterian species; whereas the “bilaterian genes” are those shared between at least two of the three bilaterian clades. The silhouettes in (B) and (C) were downloaded from PhyloPic (Nemertodermatida, Andreas Hejnol; *Chrysaora*, Levi Simons; Asteroidea, Fernando Carezzano; and *Tricolia*, Tauana Cunha).

### 2.4. Ultrafiltration excretory system

The nine genes investigated were annotated in both protostomes and deuterostomes. In Acoelomorpha, all genes but *Osr* were annotated, whereas only three out of the nine genes were found in the two cnidarian species (*ZO1*, *Six*, and *Lhx*; Fig. 3A). According to GenBank, three more genes (*Nephrin*, *Eya*, and *POU3*) are also present in this phylum (Fig. 3A).

**Figure 3:**
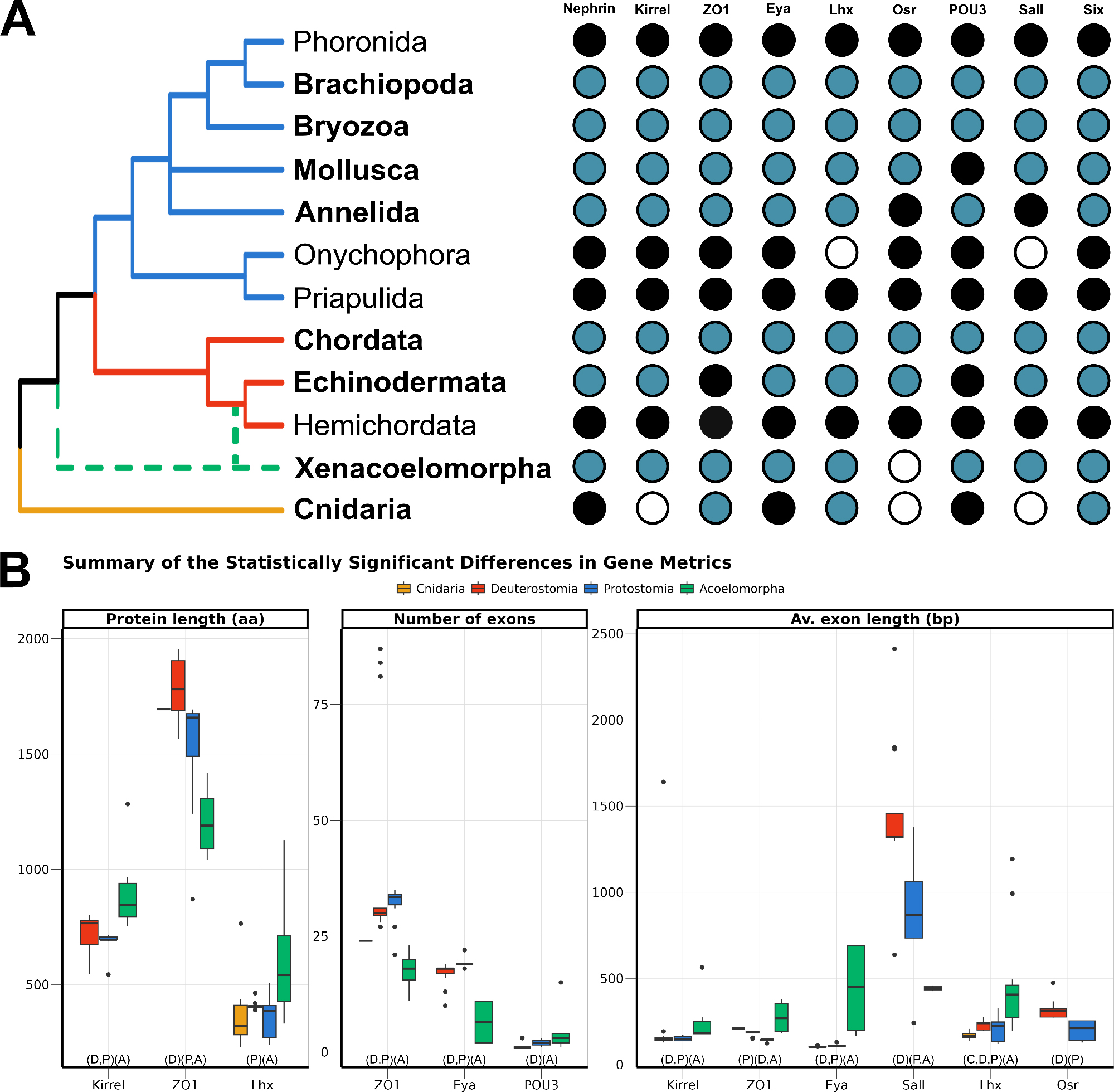
(A) Presence of the nine genes related to the ultrafiltration excretory system annotated in this study (blue), complemented with information from GenBank (black). The phyla investigated here are highlighted in bold, whereas the others were studied in Gąsiorowski et al. [20]. The cladogram topology is based on [79], including the two alternative positions of Xenacoelomorpha as a dashed line. (B) Boxplot comparing the three metrics related to gene architecture, separating the four main clades analysed per colour. Only the comparisons significantly different are shown, but the full result is included in Supplementary Figure S5. In the X-axis, below the boxplots, the brackets summarise the pairwise comparisons, clustering the clades with no significant differences within the same brackets.

The gene architecture (in terms of protein length, number of exons per gene, and average exon length) was compared for the nine genes among four clades: Cnidaria, Acoelomorpha, Deuterostomia, and Protostomia. Almost half of the 27 comparisons returned statistically significant differences among clades, most of them related to acoelomorphs (Fig. 3B). Despite the evident variation in protein length, both within and among clades, only three out of the nine genes were considered to be statistically significant: *Kirrel*, which is significantly longer in acoelomorphs; *ZO1*, longer in deuterostomes; and *Lhx*, but in this case the differences were only significant between acoelomorphs (longer) and protostomes (shorter). As for the number of exons per gene, *ZO1* and *Eya* presented fewer exons in acoelomorphs than in both deuterostomes and protostomes. Finally, the last gene with a significantly different number of exons is *POU3*. This is a relatively short protein, on average shorter than 500 amino acids in all clades, and with very few exons: only one exon in all deuterostomes but *Branchiostoma floridae* (three), between one and three in protostomes, and between one and four in acoelomorphs. Only the differences between deuterostomes and acoelomorphs were statistically significant. Two remarkable outliers were found when comparing the number of exons per gene. Three chordate *ZO1* sequences were divided into more than 80 exons (average 29.5) and one of the *POU3* sequences annotated in *P. naikaiensis* presented 15 exons (average in Acoelomorpha: 2.6). Nonetheless, these proteins were roughly of the same size as the others and their identity to the most similar protein was above 90%.

In an attempt to avoid the misleading effect of errors in the annotation (partial proteins will be generally shorter and with fewer exons), the average exon length was also considered. In this case, six out of the nine proteins were significantly different among clades. The average exon length was significantly longer in acoelomorphs in three genes (*Kirrel*, *Eya*, and *Lhx*), and two in deuterostomes (*Sall* and *Osr*, although the latter was only present in deuterostomes and protostomes). The only instance with significantly shorter exon lengths is the protostome’s *ZO1* gene. Finally, among the nine comparisons including at least one cnidarian species (three genes, three metrics) no significant differences were found but in the average exon length of *Lhx*, which is significantly shorter than that of acoelomorphs, as also observed in deuterostomes and protostomes.

## 3. Discussion

### 3.1. Performance of the Ultra-Low DNA Input protocol for sequencing large genomes

The steady development of sequencing technologies is allowing the generation of genomes spanning the diversity of life, which now includes minute organisms. Indeed, thanks to the latest low and ultra-low DNA input protocols sequencing high-quality genomes from millimetric animals is now possible [6,25,26]. In this study, we used the Pacbio Ultra-Low DNA Input protocol to sequence the genome of *N. westbladi*, reporting the first nemertodermatid genome, sequenced from a single microscopic worm. The estimated genome length is comparable to that of *P. naikaiensis*, but considerably shorter than *S. roscoffensis* and *H. miamia* [12]. Although the *P. naikaiensis* genome is slightly more contiguous than *N. westbladi*, all the metrics compared are similar between the two genomes. In contrast, both *S. roscoffensis* and *H. miamia* were scaffolded using proximity ligation data, and hence both show much higher contiguity. Beyond the differences in contiguity, annotation metrics are comparable among *N.* westbladi, *P. naikaiensis*, and *S. roscoffensis*. In this case, *N. westbladi* is more similar to *S. roscoffensis* than to *P. naikaiensis*, which shows the lowest genome completeness and number of gene models. In particular, the analysis of gene architecture shows that the number of exons per gene and intron size is also comparable, likely meaning that the annotated proteins are complete or nearly complete, facilitating the study of gene properties, such as intron-exon structure. Likewise, all genomes are similarly repetitive: *N. westbladi* 59.85%; *P. naikaiensis* 69.8%; *S. roscoffensis* 61.14%; and *H. miamia* 53%, but this is where the difference between the short- and long-read genomes of *N. westbladi* strikes the most. Although they have similar completeness and number of gene models, the Illumina genome is only 62.2 Mbps long and only 16.4% repeats, which is probably explained by the difficulty to assemble repetitive areas of the genome [27].

It is obvious from the comparisons above that achieving a highly contiguous genome from single-millimetre worms is still challenging. One potential explanation for this is the large size of acoelomorph genomes, ranging between 500 and 1100 Mbps and above the maximum genome size advised by Pacbio. The ultra-low DNA input protocol has insofar been tested in animals whose genome size ranges between 200 and 300 Mbps, returning significantly more contiguous genomes than that of *N. westbladi* [4–6]. Alternatively, the generally lower coverage of the nemertodermatid genome, due to its larger size, could have also resulted in a more fragmented assembly. Yet sequencing a second HiFi SMRT cell was not feasible due to the low DNA yield. One straightforward solution to improve genome contiguity is complementing this approach with ligation data, which has shown great results both in *S. roscoffensis* and *H. miamia* [11,12]. However, this approach would require pooling tens of individuals to obtain the required amount of DNA, which is not feasible for all animals. *N. westbladi* cannot be cultured in the lab and collecting worms in enough numbers is challenging. Interestingly, the *P. naikaiensis* genome (the most similar to *N. westbladi*) was sequenced from a pool of individuals in 52 SMRT Cells [10], whereas the *N. westbladi* genome comes from a single worm and one HiFi SMRT Cell. Altogether, these results highlight the potential of combining this protocol and HiFi to generate good-quality genomes from single, microscopic organisms, even for relatively large genomes.

The BlobTools analysis identified a high degree of contamination in the raw assembly of *N. westbladi*, which is to be expected from a microscopic organism caught in the wild. Although *N. westbladi* is known to not carry internal symbionts (based on hundreds of observations), a TEM analysis revealed the presence of gram-negative bacteria throughout the epidermal cilia [28]. Thus far, DNA extraction was performed from a whole specimen, thus sequencing the gut microbiome, and other contaminants might have been transferred from the DNA suspended in the seawater. A common practice to limit the presence of contaminants in the organism is to starve the animals before DNA extraction. Besides, the acoel genomes were sequenced from juveniles, before they incorporate the symbiotic algae, and rinsed with filtered seawater (e.g. [10,11]). However, as seen here this is not enough to prevent the presence of contaminants. This was particularly problematic in the case of *P. naikaiensis*, as almost 4% of the contigs (75 Mbps, over 10% of the genome) were identified as bacterial contigs. It is important to notice that a big fraction of the genomes did not have any hit against the Uniprot database (*N. westbladi* 13.2%; *P. naikaiensis* 8.4%; *S. roscoffensis* 1.9%; Supplementary Table S1), showing the importance of sequencing underrepresented groups to improve the reference databases.

### 3.2. Evolution of Acoelomorpha genomes

The increasing availability of animal genomes has unveiled a remarkable diversity in genome sizes, ranging from 15.3 Mbps in the orthonectid *Intoshia variabilis* to the 43 Gbps of the lungfish genome [29,30]. It has been observed that miniaturised animals tend to have smaller genomes, which has been noted both in vertebrates and invertebrates [7,9,31], but with notable exceptions to this rule, as observed in nematodes and platyhelminths [32]. Genome length in the latter ranges between 700 and 1200 Mbps, the same size range as birds, some gastropods, and many freshwater fish, among others [33–35]. Similarly, acoelomorph genomes vary between 559 and 1059 Mbps but contrast with the chromosome-level genome of *Xenoturbella bocki*, estimated at 110 Mbps [22]. Comparisons of eukaryotic genomes proposed that variations in genome sizes and proportion of repeat elements are correlated [36,37], which might also apply within Xenacoelomorpha. Acoelomorph genomes show a much higher than the small genome of *Xenoturbella* [22].

In turn, acoelomorph genomes seem to be characterised by an important reduction of gene content. Indeed, almost 60% of the genes shared between protostomes and deuterostomes are missing in acoelomorphs, which could be explained by the morphological simplicity of these worms compared with other bilaterians, but the evolutionary interpretation depends on the phylogenetic hypothesis. Under the Xenambulacraria hypothesis, their absence must be explained by massive secondary losses. The Nephrozoa hypothesis, on the other hand, suggests that the evolution of the genes exclusively shared by deuterostomes and protostomes occurred in the stem line of Nephrozoa and no *ad hoc* hypotheses of gene loss are required.

### 3.3. Evolution of the genes related to the ultrafiltration excretory system

Despite the absence of a specialised excretory system in Xenacoelomorpha, Andrikou et al. [19] described the presence of active excretion in this phylum through the digestive tissue and annotated several genes known to participate in the excretory mechanisms of nephrozoan animals. Here, we annotated in the genomes of Acoela and Nemertodermatida seven of the nine genes involved in the development of the nephridia and one more (*Sall*) in Acoela. Regardless of their phylogenetic position, whether as a sister to Ambulacraria or Nephrozoa, the presence of these genes might be explained by their participation in other important functions. A spatial transcriptomics analysis in the acoel *Isodiametra pulchra* and the nemertodermatid *Meara stichopi* located the expression of *Nephrin* in the brain and the nerve cords [19], which resembles observations in mammals and *Drosophila*, the latter through the *Nephrin* homolog *Sns* [38–40]. In contrast, no homologs to the *Osr* gene (named *Odd* in *Drosophila*) could be annotated in any of the acoelomorph genomes. A BLAST search over the two *Xenoturbella* transcriptomes failed to annotate this gene in these species, confirming its absence is a general trait of the phylum. This is noteworthy, as *Osr* is essential in the formation of the excretory organs: in vertebrates, it participates in the formation of the pronephros, the first stage in kidney formation, and its knock-out results in the absence of kidneys [41]; whereas in *Drosophila*, *Odd* participates in the embryogenesis of the tubules of Malpigi [42]. Overall, it seems that the molecular machinery that participates in the functioning of a complex ultrafiltration excretory system is present in acoelomorphs, but they lack the one gene necessary to promote the formation of discrete excretory organs.

This pattern fits well within the Nephrozoa hypothesis. In this scenario, the origin of the excretory organs would be the result of gene co-option, a common phenomenon in the origin of key innovations, such as the development of the radula and shell evolution in molluscs [43] or the multiple origins of cnidarian eyes [44]. Interestingly, six of the nine genes investigated have been annotated in different cnidarian species, strengthening the idea of the molecular machinery predating the appearance of this specialised excretory system [20]. Thus far, *Osr* has not been annotated in any phylum outside of Nephrozoa, supporting the origin of this gene in the ancestor of this clade. Nevertheless, given the ongoing debate around the phylogenetic position of xenacoelomorphs, the Xenambulacraria hypothesis also needs to be taken into consideration. If Xenacoelomorpha is the sister group of Ambulacraria, additional *ad hoc* hypotheses have to be invoked: either the *Osr* gene was independently gained in Protostomia, Ambulacraria, and Chordata or it was lost in Xenacoelomorpha. The *Drosophila Odd* gene has been shown to activate the formation of kidney tissue in vertebrates [42], which suggests a common origin of both genes in protostomes and deuterostomes. Likewise, the function of this gene is not limited to the development of the excretory organs, but it participates in the development of the foregut in vertebrates [45] and it is known to be expressed in the digestive tract of spiralians and hemichordates [20]. Although its general anatomy varies within the phylum, the presence of a sack-like gut is considered a plesiomorphy within Xenacoelomorpha [46] and the involvement of *Osr* in its development could be expected. In this light, the reduction of the excretory organs alone would not explain the secondary loss of *Osr*, as it would need to be completely nonfunctionalized before that.

We found statistically significant differences in the gene architecture of all genes but *Nephrin* and *Six*, six of them related to the average exon length. Acoelomorpha is responsible for two-thirds of the differences observed, which fits with the co-option of these genes into the development of the excretory system in the ancestor of Nephrozoa. Changes in gene structure are a strong generator of diversity, particularly after gene duplication, as part of the neofunctionalization of proteins [47]. Alternatively, the differences observed might simply be explained by changes in the selective pressures during the acquisition or the reduction of this system, something that might be supported by the observations in Bryozoa. Within protostomes, Bryozoa, which also lack an excretory system, is responsible for most of the variation observed. Notably, half of the gene metrics that are visibly different in this phylum are shared with acoelomorphs: *ZO1* and *Lhx* length, *ZO1* number of exons, and *Sall* average exon length. However, the variation does not always go in the same direction (e.g., the number of exons in *ZO1* increases in Acoelomorpha, but decreases in Bryozoa), likely because the absence of the excretory organs in the two phyla represents two independent evolutionary events. Some authors have argued that the rapid evolutionary rates observed in Acoelomorpha might be associated with other traits observed in this group, such as chromosomic rearrangements or changes in gene content, misleading comparative analyses and making *Xenoturbella* a better model for studying the evolution of Xenacoelomorpha [18,22]. Unfortunately, the genomic data of *X. bocki* is yet not available so we have inferred a gene tree for each of the nine genes analysed and compared the differences in branch lengths among clades to explore this possibility (Supplementary Figure S4). Although branch lengths are indeed significantly longer in acoelomorphs than in any other clade (except in *Lhx* and *Six*), they are also longer in deuterostomes compared to protostomes despite the similarities between the two clades. In more detail, protostomes present the shortest branches in the gene trees, while Bryozoa is one of the phyla with the most changes in gene architecture. Hence, the accelerated evolutionary rates of Acoelomorpha do not seem to be the main factor underlying the differences observed in these genes, although it would be interesting to confirm this once all the data from the *Xenoturbella* genome is publicly available.

## 4. Conclusions

In this study, we have generated the first draft of a nemertodermatid genome, sequenced from a single, microscopic individual using the Ultra-Low Input DNA protocol and HiFi. We show that this approach is capable of producing genomes of relatively good quality even from small organisms with long genomes. The main drawback is genome contiguity, which remains the main challenge and one of the avenues in genome sequencing that need the most attention. Nevertheless, genome quality is good enough to annotate full proteins, allowing detailed analysis of gene architecture. We prove this by analysing the genes related to the ultrafiltration excretory system. We observe that the molecular machinery related to this system predates its origin, as most of the genes were present in Urbilateria or even in the cnidarian-bilaterian ancestor. Interestingly, all genes but *Osr*, the one gene triggering the formation of these organs, were annotated in Xenacoelomorpha. Thus far, gene architecture is markedly different in Acoelomorpha, which cannot be explained either by the accelerated evolution of this clade or the lack of the excretory system alone. All these findings are more easily explained under the Nephrozoa hypothesis.

## 5. Material and Methods

### 5.1. DNA extractions, library preparation, and sequencing

High molecular weight DNA was extracted from single individuals of the nemertodermatid *Nemertoderma westbladi* stored in either ethanol, RNAlater, or RNA Shield using two different methods: the salting-out protocol and the QIAamp Micro DNA kit. The Qubit dsDNA HS kit, a 2% agarose gel, and a Femto Pulse system were used to ensure the extraction met the minimum requirements for DNA yield and fragment size (the majority of gDNA over 20 kbps).

Library preparation and sequencing followed the PacBio Ultra-Low DNA Input protocol with small modifications. Briefly, DNA was sheared to 10kbps using Megaruptor 3 instead of Covaris g-TUBE. After removing single-strand overhangs and repairing the fragment ends, DNA fragments were ligated to the amplification adapter and PCR amplified in two independent reactions (Reaction Mix 5A and 5B) of 15 cycles each. Amplified DNA was purified using ProNex Beads, pooled in a single sample, damage repaired for the second time, and ligated to the hairpin adapters. Size selection of the prepared SMRTbell library was done using a 35% dilution of AMPure PB beads, which removed all fragments shorter than 3kbps, instead of the BluePippin system. Finally, the library was sequenced in one SMRT cell on the Sequel IIie platform.

### 5.2. Data filtering, assembly, and decontamination

The ‘Trim gDNA Amplification Adapters’ pipeline from SMRT Link v11 was used to remove sequencing adapters. Three genome assembly strategies were attempted and compared: the IPA HiFi Genome Assembler included in SMRT Link v11 (PacBio), Hifiasm v.0.7 [48], and Flye v.2.8.3 [49]. Based on genome length, fragment size, and completeness (measured with BUSCO and the metazoa odb10 database), the Flye assembly was selected for downstream analyses, which included two additional scaffolding approaches. First, the two *N. westbladi* transcriptomes were mapped to the genome using HISAT2 v.2.0.5 [50] and fed to P_RNA_SCAFFOLDER [51]. Second, the genome of *S. roscoffensis* was used as a reference to map the assembled genome with RagTag v.2.0.1 [52]. Unfortunately, none of these attempts improved the genome contiguity any further.

The raw assembly was decontaminated following the BlobTools2 pipeline [53]. Coverage data was calculated by mapping the filtered HiFi reads to the assembled genome using Minimap2 [54], genome completeness inferred with BUSCO v.5.2.2 [55] and the Metazoa odb10 database, and taxonomic information was identified through BLAST searches of the contigs versus the UniProt database (Release 2022_05) using diamond v.0.9.26.127 [56]. Only the contigs identified as “Metazoa” were kept at this stage. Additionally, a BLAST search was used to remove mitochondrial contigs. Finally, Minimap2 was used to map the reads back to the decontaminated genome to separate the nemertodermatid reads. The k-mer approaches GenomeScope v.2.0 and SmudgePlot [57] were used to calculate the genome heterozygosity and ploidy before and after the decontamination step with a kmer length of 21. To identify the contaminant contigs, the diamond output was used to extract the *Taxid* information of the hits, which is associated with a unique taxonomic category on the NCBI database.

### 5.3. Genome annotation

RepeatMasker v.4.1.2-p1 [58] was used to soft mask the repeats in the decontaminated genome with the rmblast engine, for which a custom repeat database was generated with RepeatModeler v.2.0.1 [59] and the -LTRStruct option activated. Afterwards, the genome was annotated with BRAKER2 [60] using transcriptomic and proteomic evidence. The two available transcriptomes for *N. westbladi* were downloaded and quality filtered in a two-step approach. Adapters removal and a light trimming were performed with Trimmomatic v.0.36 (as implemented in Trinity v2.6.6, [61]), followed by a more thorough cleaning with PRINSEQ v.0.20.3 [62]: trim all terminal bases with a quality below 30 and filter out reads whose mean quality is below 25, low complexity sequences (minimum entropy 50), and reads shorter than 75bp. Clean reads were mapped to the soft-masked genome with STAR v.2.7.9 [63] and the options “--sjdbOverhang 100 --genomeSAindexNbases 13 --genomeChrBinNbits 15” and “--chimSegmentMin 40 --twopassMode Basic”. For the proteomes, the gene models from the acoel *P. naikaiensis* [10], the BUSCO Metazoa odb10 database, and a custom set of single-copy orthogroups, inferred from published transcriptomes with OrthoFinder v.2.4.1 [64], were concatenated and mapped to the *N. westbladi* genome using ProtHint v.2.6 [65]. The inferred gene models were functionally annotated by pfam_scan v.1.6 [66] and the PFAM 31.0 database.

### 5.4. Quality control

The quality of the decontaminated genome was assessed using QUAST v.5.2.0 [67] and the completeness of the genome and the annotation with BUSCO v.5.2.2 using the Metazoa and Eukaryota odb10 databases. Since all the metazoan contigs were kept during the decontamination step, two approaches were followed to ensure they belong to the nemertodermatid genome. First, a distance tree was inferred with FastMe v.2.1.5 [68] based on a distance matrix calculated with Skmer [69], an alignment-free method designed to estimate genomic distances, over the *N. westbladi* genome and 18 metazoan genomes downloaded from GenBank (Supplementary Table S4). Second, a phylogenetic tree was inferred from these genomes except for three for which the annotated proteome was not available. Briefly, orthogroups were inferred with OrthoFinder v.2.4.1 [64] and clean from paralogs with PhyloPyPruner v.1.2.3 [70] using the “Largest Subtree” method, collapsing nodes with bootstrap support lower than 60, and pruning branches more than five times longer than the standard deviation of all branch lengths in the tree. Then, orthogroups were aligned with MAFFT v.7.475 using the L-INS-i algorithm [71], cleaned from poorly aligned sites with BMGE v.1.12 [72], tested for stationarity and homogeneity (symmetry tests) with IQ-TREE2 v.2.1.3 [73], and concatenated with FASconCAT v.1.05 [74]. Finally, a phylogenetic tree was inferred using coalescence (ASTRAL; [75]) and site-specific, concatenation-based methods (assuming 20 amino acid categories, C20) with IQ-TREE v.1.6.12 [76].

All the genome metrics, including length, contiguity, number of genes, and completeness, among others, were compared to the acoel genomes from *P. naikaiensis* [10] and *S. roscoffensis* [11], which were also tested for contaminants using BlobTools2, following the same pipeline and with the same filtering criteria. The genomes of *Hofstenia miamia* and *Xenoturbella bocki* [12,22] were not considered because an annotation file with details of protein structure is not available for any of them. Additionally, a second *N. westbladi* genome sequenced in an Illumina HiSeq2500 platform was also included in the comparisons to estimate the improvement in genome quality with HiFi data from a short-read approach. Briefly, DNA was extracted from a pool of 12 individuals, collected in the same location at the same time, the sequencing library was prepared with a Rubicon kit, and the sequencing generated more than 385 million reads. The Illumina reads were assembled with SPAdes v.3.14.1 [77], with four kmer lengths (21, 33, 55, 75) and error correction activated. Finally, this genome was analysed with the same parameters as the HiFi genome to eliminate contamination contigs, produce completeness stats, and annotate gene models.

### 5.5. Analysis of gene content

To analyse the evolution of gene content in Acoelomorpha, the annotated genomes of 18 animals were compared, including *N. westbladi* (Nemertodermatida) and *P. naikaiensis* and *S. symsagittifera* (Acoela) as representatives of Acoelomorpha, eight protostome genomes, four deuterostomes, and three cnidarians as the outgroup to Bilateria (Supplementary Table S4). Redundancies in the gene models of all genomes were removed with CD-HIT [78], clustering all sequences more than 95% identical, and then functionally annotated with pfam_scan v.1.6 [66] and the PFAM 31.0 database. The annotated proteins were clustered using OrthoFinder v.2.4.1 [64] and used to calculate the number of genes specific to or shared among the four main clades of interest: Cnidaria, Acoelomorpha, Deuterostomia, and Protostomia. The genes present in at least one cnidarian and one bilaterian were considered to be shared across Metazoa, whereas the genes present in at least two of Acoelomorpha, Deuterostomia, and Protostomia were considered to be shared across Bilateria. Then, the proportion of “metazoan” and “bilaterian” genes absent from each of the three bilaterian clades was calculated based on these two datasets.

### 5.6. Annotation and comparison of the genes related to the ultrafiltration excretory system

This analysis was based on the results of Gąsiorowski et al. [20], who used spatial transcriptomics to identify the genes involved in the development of the ultrafiltration excretory system in several protostomes and one hemichordate species. All the protein sequences annotated in this study were downloaded from GenBank except *Hunchback*, as they found no evidence of this gene being involved in nephridiogenesis, for a total of three structural proteins: *Nephrin*, *Kirrel*, and *ZO1*; and six transcription factors: *Eya*, *Lhx1/5*, *Osr*, *POU3*, *Sall*, and *Six1*. These genes were annotated in the same genomes used to analyse gene content evolution through BLAST searches with diamond v0.9.26.127 [56]. The correct identification of these genes was later confirmed through phylogenetic analyses with IQ-TREE v.1.6.12 [76] and manual BLAST searches on the NCBI webserver. The identification of the *Lhx1/5* and *Six1* transcription factors was not always straightforward, as they are thoroughly mixed in the phylogenetic tree with many other gene variants and sometimes different isoform names were proposed in the BLAST searches for the same sequence, and thus they represent a mixture of isoforms of the same gene. A custom R script was written to locate the filtered genes in the GFF files and extract three metrics related to gene architecture: protein length, number of exons per protein, and average exon length per gene. Unfortunately, the GFF annotation file was not available for all these genomes, so not all of them could be included in this analysis (Supplementary Table S4). To ameliorate the misleading effect of highly fragmented genes we filtered out all proteins shorter than half of the average protein length of the respective gene (a total of 10 proteins). To test if the observed differences in the three gene metrics were statistically significant, the Shapiro-Wilk’s method and the Barlett test were used to check if they follow a normal distribution and the homogeneity of their variances, respectively. For each gene, the differences among clades were tested with either an ANOVA or a Kruskal-Wallis test, depending on the result of the normality and homoscedasticity tests. Finally, the Bonferroni correction (ANOVA) and the Dunn test (Kruskal-Wallis) were selected to run pairwise comparisons in all cases identified as statistically different.

## 6. Data availability

The raw sequencing data and the annotated genome assemblies are available through the NCBI database under BioProject PRJNA981986. Raw and decontaminated assemblies, as well as annotation files, predicted nucleotide and protein sequences, mapped reads, and supporting information were deposited in the GigaScience database GigaDB. The code necessary to replicate all the analyses has been uploaded to the GitHub repository https://github.com/saabalde/2023_Nemertoderma_westbladi_genome

## 7. Additional files

**Supplementary Figure S1:** Summary of the completeness analyses performed after the decontamination. The four genomes were analysed with BUSCO using the Eukaryota (A) and Metazoa (B) odb10 databases.

**Supplementary Figure S2:** Ploidy result generated by SmudgePlot after the decontamination (kmer = 21).

**Supplementary Figure S3:** Transformed plot generated by GenomeScope analysis after decontamination (kmer = 21).

**Supplementary Figure S4:** Average branch length per clade and ultrafiltration gene. The error bars represent the standard error.

**Supplementary Figure S5:** Summary of the analyses related to the evolution of the ultrafiltration excretory system. (A) Phylogenetic tree inferred with IQ-TREE to confirm the correct annotation and monophyly of the genes. Boxplot summarising the (B) protein length, (C) number of exons per gene, and (D) average exon length per clade and gene. The results are presented as a facet to separate the structural proteins and transcription factors in two panels. For the two panels, the same scale in the Y-axis is used.

**Supplementary Table S1:** Summary of the contaminants identified in the *N. westbladi* genome by BlobTools2.

**Supplementary Table S2:** Statistics of the repeat elements identified and masked by RepeatMaster. The abundance of each repeat family is shown as a percentage of the genome length.

**Supplementary Table S3:** List including the taxonomic information, to the lowest category possible, of all the contaminants identified in the assembly of the *N. westbladi* genome (HiFi).

**Supplementary Table S4:** Accession number and reference of the genomes downloaded from the SRA and used in comparative analyses.

## 8. Acknowledgements

We are thankful to C. Laumer for his advice during the early stages of this project. Analyses and data handling were enabled by resources in projects SNIC 2020/15-191 and SNIC 2021/22-562 provided by the National Academic Infrastructure for Supercomputing in Sweden (NAISS) at UPPMAX, funded by the Swedish Research Council through grant agreement no. 2018-05191.

## 11. Authors’ contribution

SA, OVP, and UJ conceived the project; SA performed DNA extractions; JH was responsible for library preparations and sequencing; CTR carried out the post-sequencing analyses, from quality filtering to genome assembly; SA decontaminated and annotated the genome and performed comparative analyses; SA and UJ led the writing of the manuscript. All authors read and approved the final manuscript for submission.

