## Supplementary figures and images for "The draft genome of the microscopic *Nemertoderma westbladi* sheds light on the evolution of Acoelomorpha genomes"

### SuppMatFig_S1-BUSCO_results.png

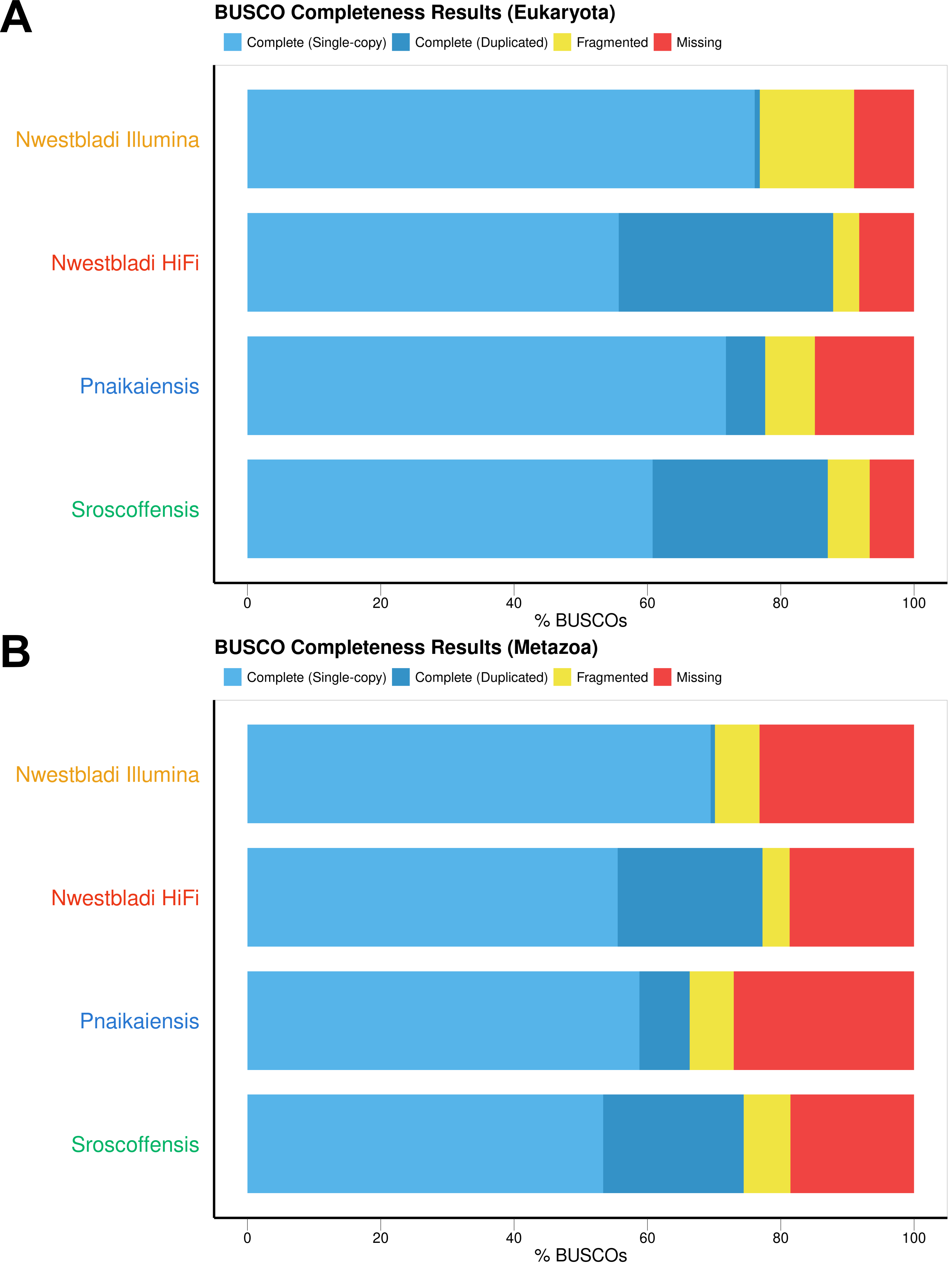

### SuppMatFig_S2-SmudgePlot.png

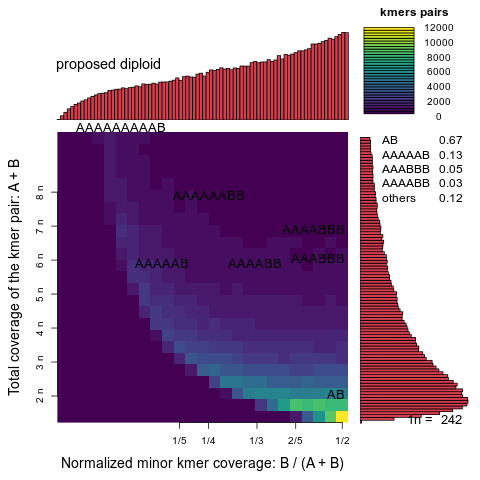

### SuppMatFig_S3-GenomeScope.png

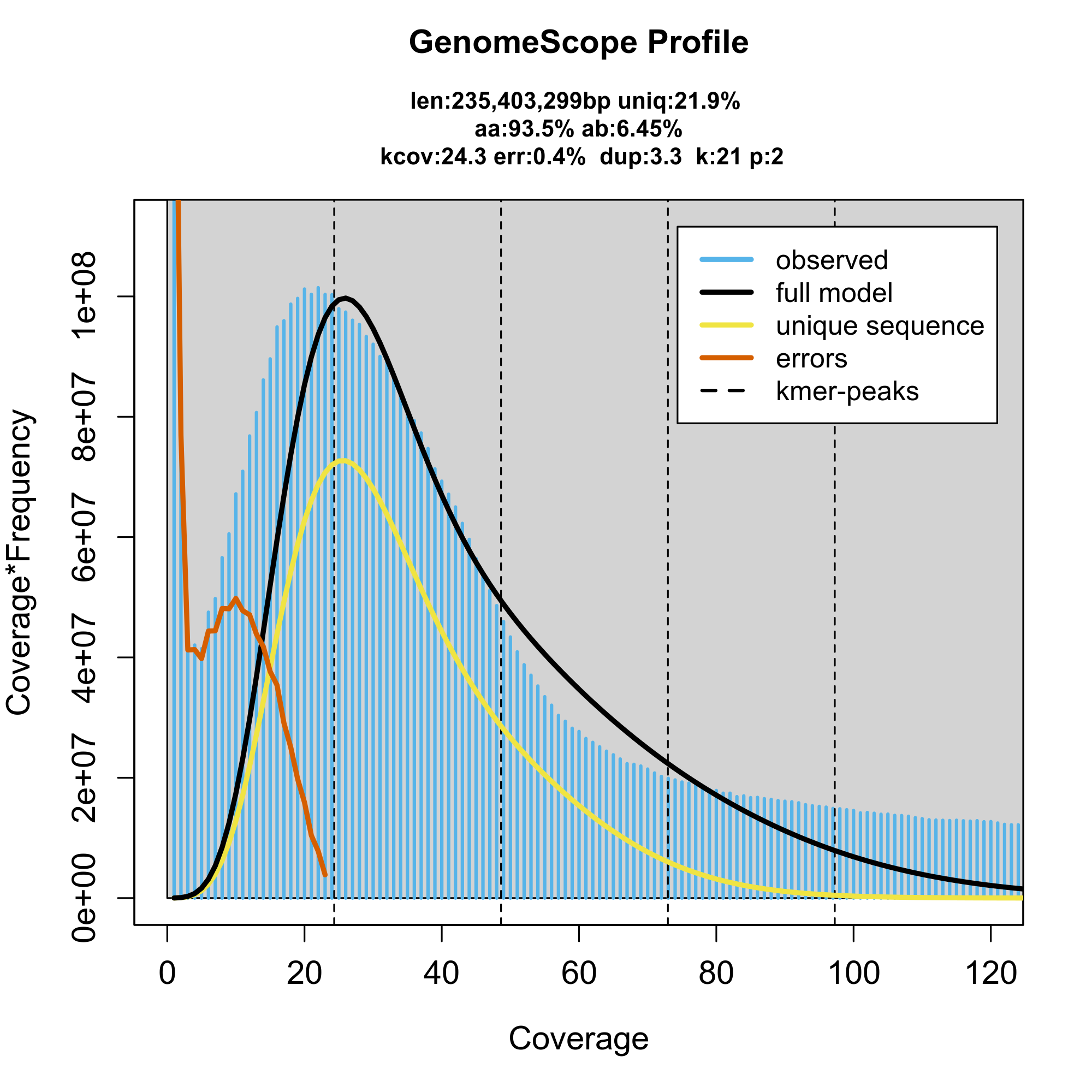

### SuppMatFig_S4-SummaryUltrafiltrationGenes.BranchLengths.png

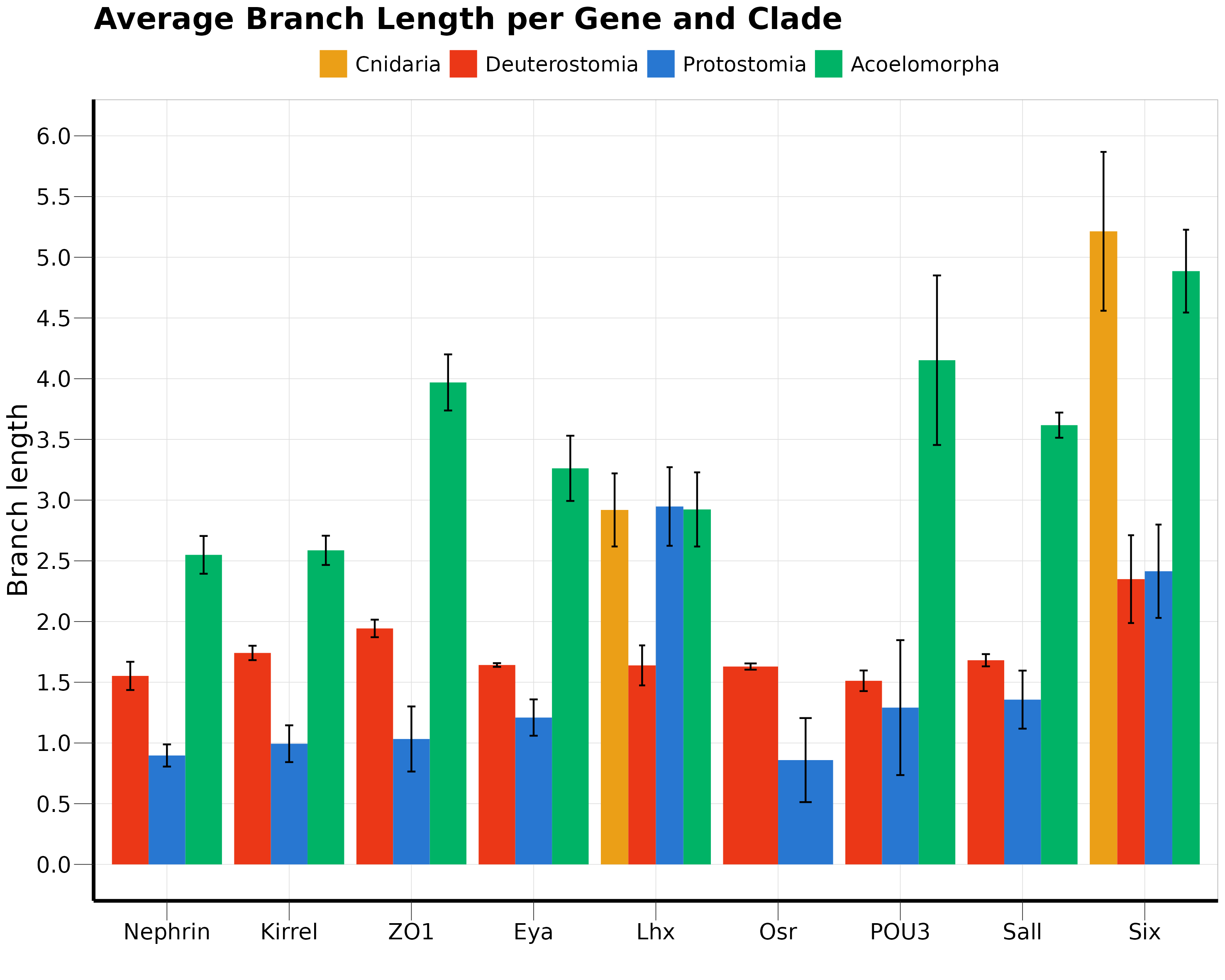

### SuppMatFig_S5-SummaryUltrafiltrationGenes.png

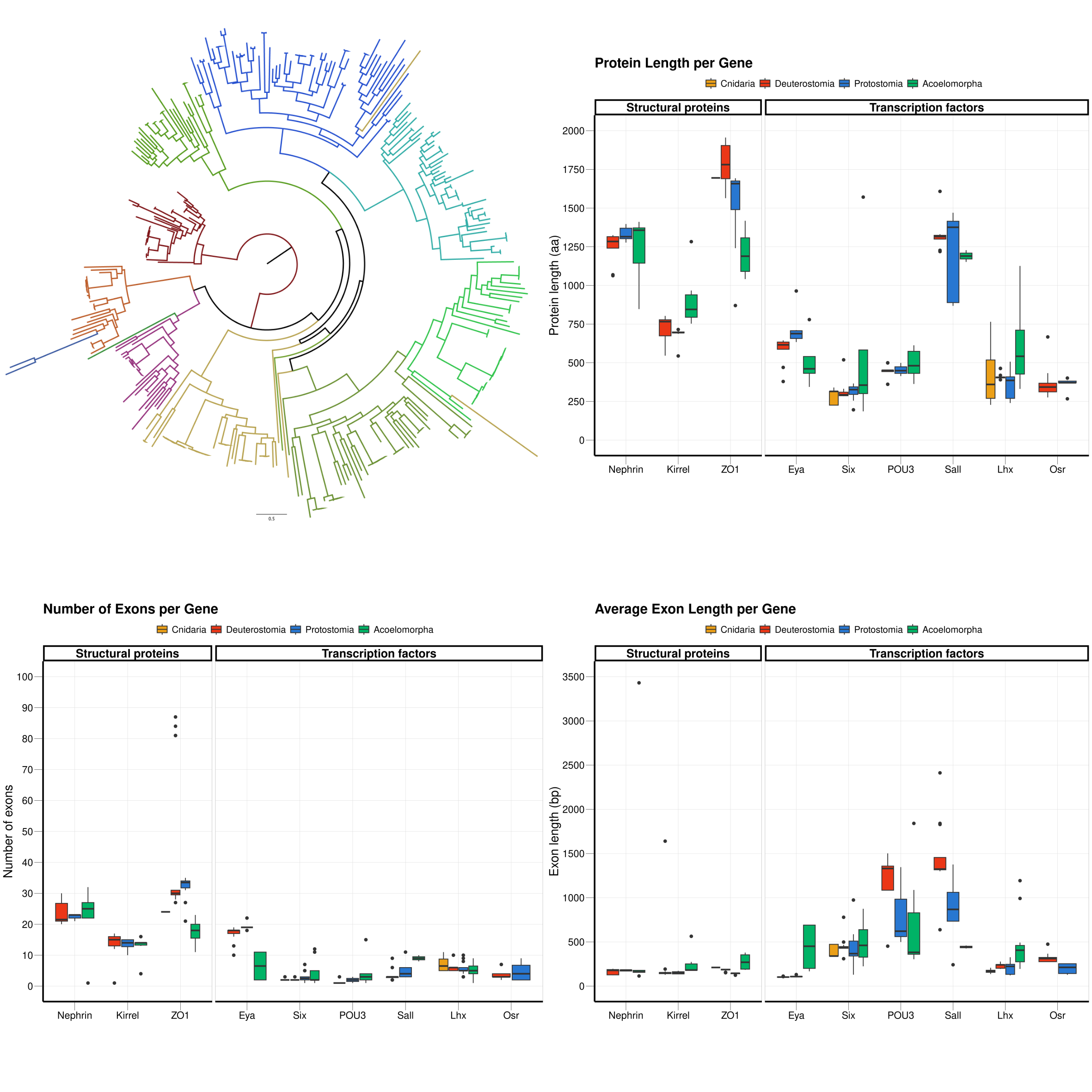
